# ToxinPred 3.0: An improved method for predicting the toxicity of peptides

**DOI:** 10.1101/2023.08.11.552911

**Authors:** Anand Singh Rathore, Akanksha Arora, Shubham Choudhury, Purva Tijare, Gajendra P. S. Raghava

## Abstract

Toxicity emerges as a prominent challenge in the design of therapeutic peptides, causing the failure of numerous peptides during clinical trials. In 2013, our group developed ToxinPred, a computational method that has been extensively adopted by the scientific community for predicting peptide toxicity. In this paper, we propose a refined variant of ToxinPred that showcases improved reliability and accuracy in predicting peptide toxicity. Initially, we used BLAST for alignment-based toxicity prediction, yet coverage was limited. We adopted a motif-based approach with MERCI software to identify unique toxic patterns. Despite specificity gains, sensitivity was compromised. We developed alignment-free methods using machine/deep learning, achieving a balance sensitivity and specificity of prediction. A deep learning model (ANN – LSTM with fixed sequence length) developed using one-hot encoding attained a 0.93 AUROC and 0.71 MCC on independent data. The machine learning model (extra tree) developed using compositional features of peptides achieved 0.95 AUROC and 0.78 MCC. Lastly, we developed hybrid or ensemble methods combining two or more models to enhance performance. Hybrid approaches, including motif-based and machine learning, achieved a 0.98 AUROC and 0.81 MCC. Evaluation on independent data demonstrated our method’s superiority. To cater to the needs of the scientific community, we have developed a standalone software, pip package and web-based server ToxinPred3 (https://github.com/raghavagps/toxinpred3 and https://webs.iiitd.edu.in/raghava/toxinpred3/).

**Author’s Biography:**

1. Anand Singh Rathore is currently pursuing a Ph.D. in Computational Biology at the Department of Computational Biology, Indraprastha Institute of Information Technology, New Delhi, India.
2. Akanksha Arora is currently pursuing a Ph.D. in Computational Biology at the Department of Computational Biology, Indraprastha Institute of Information Technology, New Delhi, India.
3. Shubham Choudhury is currently pursuing a Ph.D. in Computational Biology at the Department of Computational Biology, Indraprastha Institute of Information Technology, New Delhi, India.
4. Purava Tijare is a Project Fellow in Computational Biology at the Department of Computational Biology, Indraprastha Institute of Information Technology, New Delhi, India.
5. Gajendra P. S. Raghava is currently working as a Professor and Head of the Department of Computational Biology, Indraprastha Institute of Information Technology, New Delhi, India.

**Highlights:**

- Implementation of alignment or similarly based techniques for predicting toxic peptides.
- Discovery of toxicity-associated patterns and identification of toxic regions in peptides.
- Development of machine and deep learning-based models for toxicity prediction.
- Ensemble methods that combine alignment-based and alignment-free methods.
- Web server and standalone software package for screening toxicity in peptides/proteins.

## Introduction

Proteins are among the basic biological molecules that occur naturally in living organisms and play important roles in a number of physiological processes, such as programmed cell death and regulation of gene expression [1]. Proteins and peptides are important for maintaining body tissues and organs and are used in the diagnosis and treatment of diseases, and their stability, toxicity, and immunogenicity are primary concerns when using them as therapeutics [2]. Toxicity testing of therapeutic lead peptide molecules is critical in peptide-based drug development. The toxic properties of unidentified proteins and peptides can be evaluated by experimental toxicological approaches such as in vivo and in vitro techniques. However, timely risk assessment, assessment costs, and the need to reduce animal testing are the main reasons why in silico techniques such as molecular modeling and machine learning (ML) are preferred in the predictive characterization of protein toxicity [3]. Modeling and predicting therapeutic peptides with desirable physicochemical properties is now easier with a variety of in silico techniques for predicting protein and peptide toxicity and developing protein therapeutics.

Computational methods for molecular toxicity detection have attracted increased attention and undergone rapid advancement due to development in the field of machine learning, deep learning, and its effective use in many areas of bioinformatics, including prediction of drug-target interactions. There are two main steps involved in these computational approaches to predict toxicity. First, a number of feature extraction techniques are used to extract protein features from the source sequences. The classification process is the next stage, where the collected features are sent to a classifier to create data labels (e.g., positive or negative) [1].

Many chemical toxicity studies have been published over the years, including DeepTox [4], ProTox-II [5], and eToxPred [6]. The ability to predict the toxicity of proteins and peptides has been explored to a limited extent. For example, in 2007, prediction algorithms called BTXpred [7] and NTXpred [8] were developed to classify bacterial toxins and neurotoxins, respectively. Most tools available today are highly specialized in removing toxins from specific animal sources. A classification of animal toxins based on their major protein sequences was proposed by Naamati et al. is ClanTox [9]. Another machine learning based technique called SpiderP was developed to predict propeptide cleavage sites in spider venoms [10]. Gupta et al. published a general technique called ToxinPred in 2013 to predict the toxicity of peptides regardless of their source [11]. The scientific community makes extensive use of this technique to predict peptide toxicity. It is an SVM-based technique that looks for toxic motifs or regions formed by sequences based on variables, including amino acid composition (AAC) and dipeptide composition (DPC). Similarly, Gaces et al. created the ToxClassifier in 2016 to detect venom toxins from other proteins [12]. ToxDL [13] and TOXIFY [12] are two deep learning-based techniques that hit the market in 2019 and 2020, respectively. With TOXIFY, animal toxic proteins can be distinguished from non-toxic proteins, and the toxicity of animal proteins can be determined with ToxDL. A machine-learning technique called NNTox was developed in 2019 to identify protein toxicity based on several gene ontology annotations [14]. Wei et al. developed deep learning-based methods such as ATSE [15] and ToxIBLT [16] to predict protein/peptide toxicity based on structural, evolutionary, and physicochemical sequence features. ToxinPred2 is another recently developed machine learning based tool to classify the toxic and non-toxic protein sequences, which is trained and validated on experimentally validated large protein toxins [17]. To Enhance Protein and Peptide Toxicity Prediction, Zhao et al. published Channel Attention-Integrated Convolutional Neural Network and Gated Recurrent Units (GRU) based method [18]. A novel end-to-end deep learning-based method called ToxMVA was developed by Hua Shi et al. to predict protein toxicity [1]. To predict protein toxicity, a new technique called CSM toxin [19]was developed. It is based on the ProteinBERT deep learning model. ProteinBERT is comparable to the natural language processing paradigm BERT (Bidirectional Encoder Representations of Transformers) [20].

In this work, we attempted to establish a highly robust approach to predict peptide toxicity that would enhance our previous method, ToxinPred. While ToxinPred is highly accurate and popular, several limitations still hinder its development. One major issue with ToxinPred was that it was trained on only 1805 toxic peptides. We have proposed an improved technique called “ToxinPred3.0” that is trained and evaluated on a large data set of toxin peptides to classify toxic and non-toxic peptide sequences. The models created in this work were trained and tested using the most recent dataset, which included 5,518 toxic sequences. In addition, ToxinPred3.0 has various improvements that enhance the performance of the high-accuracy model.

## Material and Methods

### Creation and collection of the dataset

Previous studies have shown that a robust dataset is necessary to train a prediction model with good generalization properties [21]. A new peptide toxicity dataset is constructed in this study. We have collected short sequences of toxic proteins or peptides from different databases and studies that include Conoserver [22], DRAMP [23], CAMPR3 [24], dbAMP2.0 [25], YADAMP [26], DBAASP-v3 [27] and UniProt release 2021_03 (released on 2 June 2021)[28]. In our study, we excluded proteins/peptides that exceeded 35 residues or contained non-natural amino acids. However, there is a chance that the SwissProt [29] database may still contain a number of toxic peptides obtained from different sources. Therefore, we removed any identical toxic proteins/peptides from the data. This led us to a set of 5518 unique toxic proteins/peptides. We also searched for toxic proteins/peptides in SwissProt using specific criteria such as the keywords “toxin” or “toxic” and limiting the search to reviewed entries. These proteins/peptides were regarded as positive datasets for toxic peptides. While retrieving well-annotated or experimentally validated toxic peptides is feasible, obtaining non-toxic ones is challenging. Therefore, we searched for protein/peptide sequences in UniProt to construct a negative or non-toxic dataset using the keywords ‘NOT toxin NOT toxic AND reviewed: yes’. In our data extraction process, we specifically targeted protein or peptide sequences that were shorter than 35 amino acids in length and eliminated any sequences containing non-natural amino acids or identical sequences. By excluding non-natural amino acids, we focused solely on naturally occurring protein or peptide sequences, which are more representative of biological systems. This resulted in a set of 4321 non-toxic peptides as negative datasets. It is important to ensure that the data used to train a toxicity detection model is balanced in terms of the number of positive and negative examples. This is because a model that is trained on unbalanced data is more likely to overfit or underfit the data [30]. We randomly generated peptides from the SwissProt database to ensure a balanced representation of positive and negative datasets. Ultimately, we arrived at 5518 unique positive and negative datasets. The creation of datasets is depicted in Figure 1.

**Figure 1:**
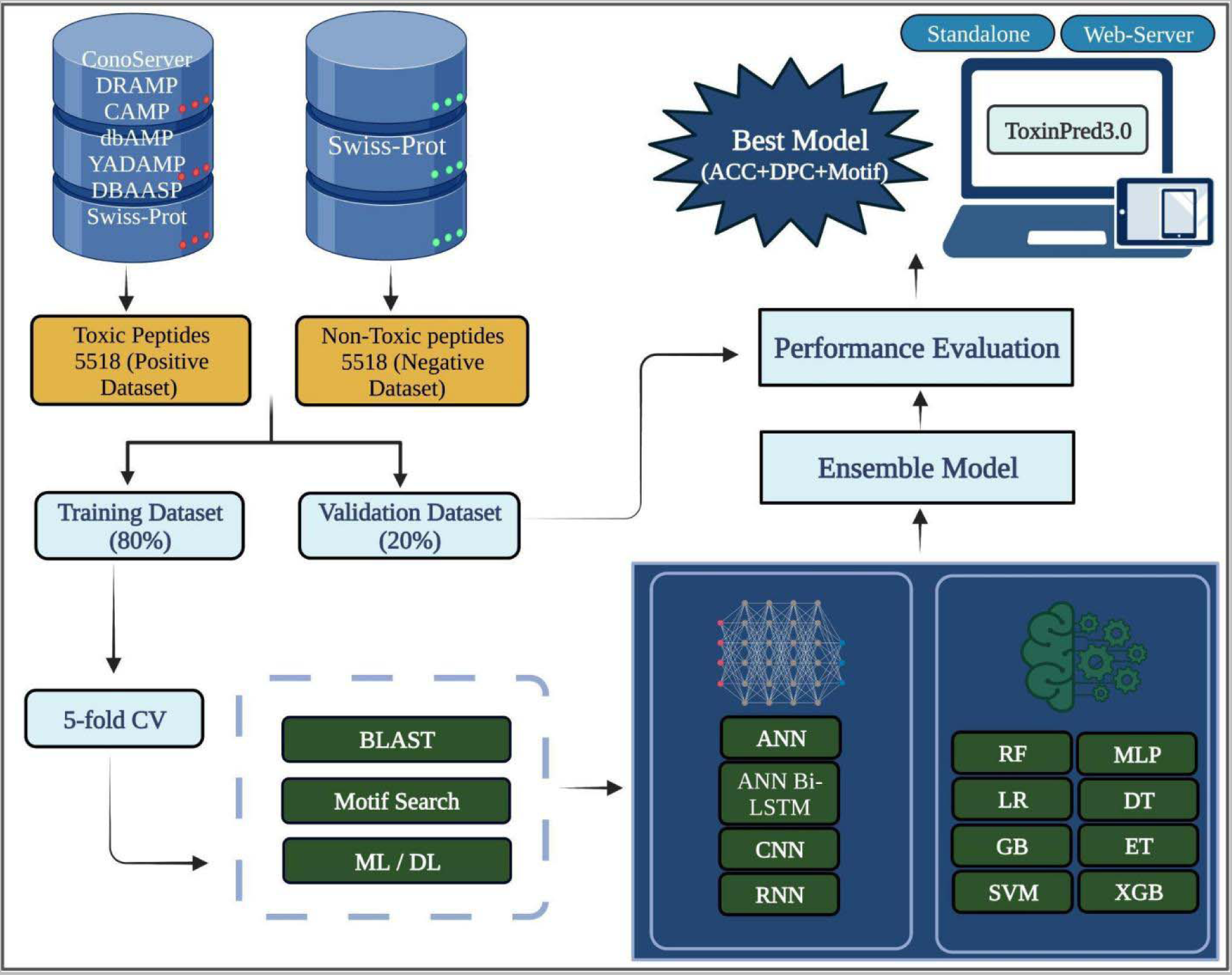
ToxinPred3.0 is depicted in the flowchart.

### BLAST search

Basic Local Sequence Alignment Tool (BLAST), version 2.9.0, is used in this study to identify toxic peptides. This tool is often employed to discover and annotate nucleotide and protein sequences [31]. We have specifically used BLASTshortP, a method designed for identifying short proteins, i.e., peptides, based on the resemblance between the sequences. To accomplish our objective, our initial step involved constructing a repository consisting of both toxic and non-toxic peptides. Once the repository was established, we proceeded to hit the query sequences against this repository to categorize them as toxic or non-toxic. This procedure was performed for various e-value cut-offs ranging from 10^−6^ to 10^2^. The repository was established by incorporating all the sequences present in the training dataset. To assess the performance for the training dataset, we took the first hit after the self-hit into account. On the other hand, when evaluating the independent dataset, the first hit was taken to establish whether a sequence is toxic or non-toxic. For example, if the query sequence hit a toxic peptide, the sequence was categorized as toxic, and similarly, for non-toxic peptides.

### Motif analysis

The Motif-EmeRging and with Classes-Identification (MERCI) tool was used to identify recurring patterns, or motifs, in toxic proteins [31,32]. Motifs are short amino acid sequences often found in proteins that perform a similar function. The MERCI tool uses a Perl script to search for motifs in a set of sequences using default parameters. This analysis provides information about the amino acid sequences that are associated with toxicity. The MERCI tool was used to analyze a set of 5518 toxic proteins. The analysis identified a number of motifs that were common to these proteins. These motifs were then used to create a predictive model that could be used to identify new toxic proteins. The identification of motifs in toxic proteins is a valuable tool for understanding the molecular basis of toxicity. This information can be used to develop new drugs and therapies to treat toxic diseases. The MERCI tool has been applied extensively to detect motifs in various toxic proteins, including proteins involved in cancer, neurodegenerative diseases, and infectious diseases [32].

### Deep learning (DL) based classifiers

During our research, we implemented distinct deep learning architectures using the Keras framework to address the classification task. The deep learning architecture offers various approaches for modeling sequential data and understanding its nature. The first model, an Artificial Neural Network (ANN) with fixed sequence length LSTM layers, efficiently captures long-term dependencies in sequential data by considering a consistent sequence length throughout the dataset. In contrast, the second model, an ANN with LSTM layers and variable sequence length, is adaptable to handle variable-length sequential data, making it well-suited for real-world scenarios where sequence lengths vary. The third model combines an ANN architecture with Bidirectional LSTM (Bi-LSTM) layers, enabling the model to process input sequences in both forward and backward directions. This enhances the ability of the model to comprehend temporal dependencies within the data. The fourth model, a Recurrent Neural Network (RNN) with LSTM layers, is specifically designed to handle sequential data by retaining memory of past time steps, making it effective for tasks such as natural language processing and time series analysis. Finally, the fifth and last model integrates a Convolutional Neural Network (CNN) with Regularized Dense layers that employ L2 regularization. This architecture is particularly useful for structured data or sentiment analysis, as the CNN component extracts local patterns while regularization techniques improve generalization and mitigate overfitting.

The main distinction among these architectures is how they handle sequence lengths. Fixed sequence lengths impose a uniform structure on all the sequences, simplifying the model architecture but potentially sacrificing information. On the other hand, variable sequence lengths with post-padding and post-truncation retain the original sequence lengths, preserving the information but requiring additional computational handling. After carefully examining the behavior of all, we have used the best performing, balanced model to be incorporated in the standalone to accurately predict the toxic and non-toxic peptides.

### FEATURE GENERATION

We use the Pfeature [33] standalone package to generate features of datasets. We considered all composition-based features and calculated a vector consisting of 9163 features for each sequence in the dataset. Supplementary Table S1 tabulates the information for each feature and the vector length.

### Feature Selection and Ranking

Previous studies have shown that not all features carry equal significance. Consequently, one of the primary obstacles lies in effectively identifying and selecting the most relevant features from a vast pool of features. With this in mind, our study has delved into various feature selection techniques, including support vector classifier (SVC) with L1-based feature selection technique (SVC-L1), Tree-based feature selection, recursive feature elimination (RFE), SelectKBest (k-best), variance threshold (VTh), and minimum Redundancy Maximum Relevance (mRMR) to address this challenge comprehensively. By implementing these methods, we aim to select the most significant set of features from a vast pool of 9163 features that bear the utmost relevance to our investigation. The top-ranked features obtained from each method are shown in Supplementary Table S2. However, when it comes to creating various machine learning prediction models, their feature selection technique performance is nearly on par with AAC and DPC - based machine learning prediction models.

### Machine learning (ML) based classifiers

A variety of machine learning techniques have been employed to differentiate between toxic and non-toxic peptides. In order to construct classification models, the following machine learning algorithms were utilized: Extra Tree (ET), Random Forest (RF), Multi-layer Perceptron Classifier (MLPC), Logistic Regression (LR), XGBoost (XGB), Decision Tree (DT), and Support Vector Machine (SVM). These classifiers underwent optimization by exploring a range of hyperparameters, and the most favorable outcomes were selected for integration into the final models. The comprehensive workflow of ToxinPred3.0, including the implementation, is visually represented in the accompanying Figure 1.

### Cross-validation and performance metrics

The datasets used in this study were divided 80:20, with 80% (i.e., 4414 toxic and 4414 non-toxic peptides) used for training and 20% (i.e., 1104 toxic and 1104 non-toxic peptides) for validation. Stratified 5-fold cross-validation was used in this instance to assess the machine learning models using 80% of the training data. While stratified 5-fold cross-validation shares similarities with k-fold cross-validation alone, it differentiates itself by incorporating a stratified sampling approach rather than a random selection. As a result, stratified 5-fold cross-validation ensures that the distribution of samples across different classes or categories remains proportional and representative within each fold. The training data is split into five portions for the internal validation, four of which are used for training and the fifth for testing. Five iterations of the procedure are carried out in order to test each of the five-folds at least once. The performance of various machine learning models was assessed by employing well-established evaluation criteria, such as threshold dependent and independent metrics. Sensitivity, specificity, accuracy, and the Matthews correlation coefficient (MCC) are under the category of threshold-dependent metrics, as their values are influenced by the classification threshold used. On the other hand, the area under the receiver operating characteristic curve (AUC) remains independent of the threshold and provides a comprehensive measure of a model’s discriminative ability. These evaluation standards have undergone extensive review and annotation in prior investigations, establishing their reliability and importance in evaluating model performance.

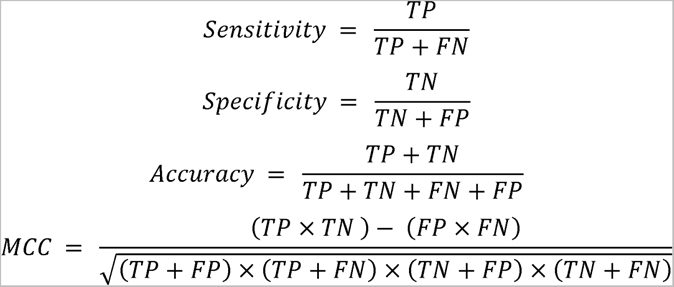

where *FP*, *FN*, *TP,* and *TN* are false positive, false negative, true positive, and true negative, respectively.

### Ensemble approach for classification

In this study, we have incorporated an ensemble/hybrid approach to improve the predictive capabilities of our final model. This hybrid approach involves a weighted scoring method, combining two distinct methods: (i) A motif-based approach utilizing MERCI and (ii) deep learning/ machine learning based technique. To begin with, we employed the MERCI motif-based approach to classify the given peptide sequence. Each prediction received a weight of ‘+0.5’ for positive predictions (indicating toxic peptides), ‘−0.5’ for negative predictions (indicating non-toxic peptides), and ‘0’ for cases where no matches were found. This weight assignment provides a quantitative measure of confidence in each prediction. In the hybrid approach, the scores obtained from both the MERCI method and the machine learning or deep learning scores were combined to calculate an overall score. Considering various threshold values, the peptide sequence could be classified as toxic or non-toxic based on this overall score. This hybrid methodology has been widely employed in numerous research studies, demonstrating its effectiveness in improving prediction accuracy. By combining the strengths of motif-based and machine learning/ deep learning techniques, our hybrid approaches (MERCI+ML and MERCI+DL) provide a comprehensive and robust framework for peptide classification, enabling more accurate toxicity predictions.

## Results

In this study, we employed a total of 5518 toxic peptides as a positive dataset and 5518 non-toxic peptides as a negative dataset. All the subsequent analyses and predictions are performed on toxic and non-toxic peptides.

### Compositional analysis

This investigation involved the calculation of amino acid composition (AAC) for both toxic and non-toxic peptides. Our analysis revealed that the average AAC of amino acid residues such as cysteine, histidine, proline, and asparagine exhibited a higher prevalence in the toxic sequences. In contrast, valine, threonine, arginine, glutamine, methionine, leucine, isoleucine, phenylalanine, and alanine are higher in the non-toxic sequences. We also compared the average AAC between the peptides and proteins of ToxinPred and ToxinPred3.0. Notably, we observed a remarkable abundance of cysteine and proline residues in the peptides derived from ToxinPred. In ToxinPred3.0, we find the same trend in composition as ToxinPred. The comparison of average AAC between ToxinPred and ToxinPred3.0 is depicted in Figure 2.

**Figure 2:**
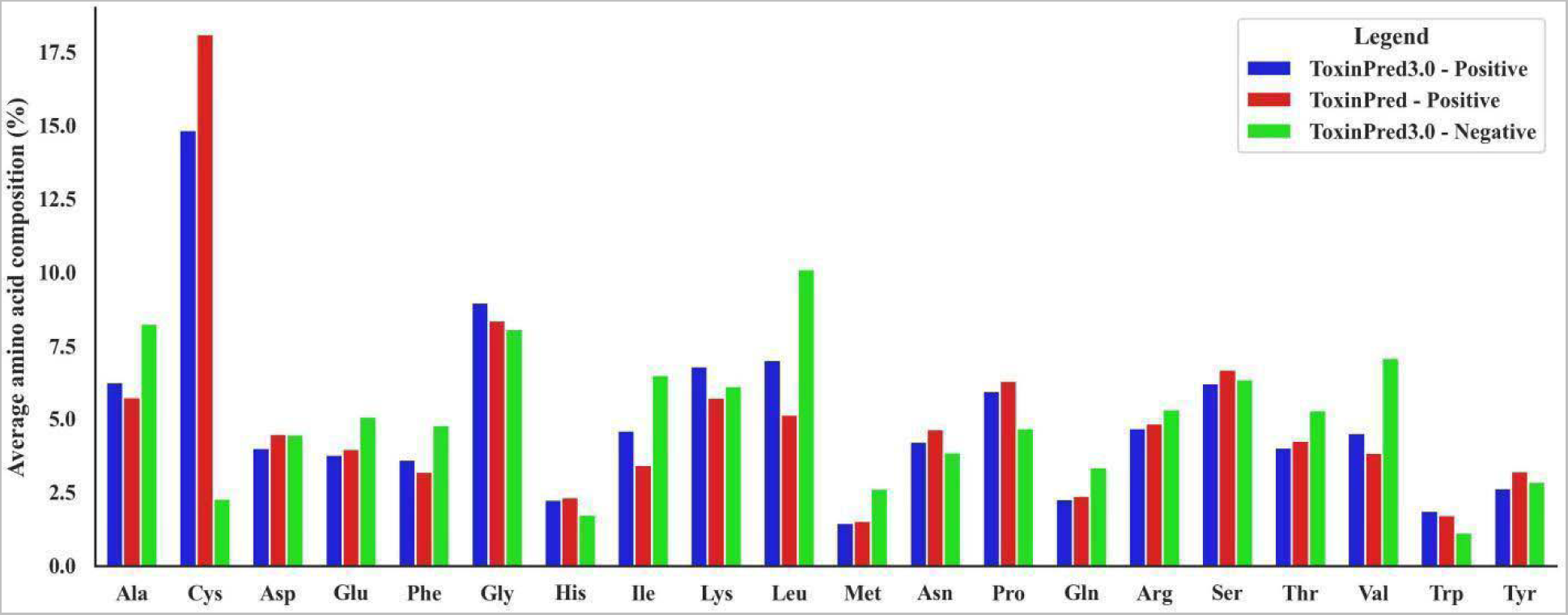
Comparison of average amino acid composition (AAC) between ToxinPred and ToxinPred3.0.

### Positional analysis

During this analysis, our focus is on investigating the preference of specific amino acids at particular positions within the peptide sequence. To accomplish this, we construct a Two logo sample (TSL) for both toxic (positive) and non-toxic (negative) peptides, as depicted in the Figure 3. The TSL provides valuable information regarding the proportional representation or relative abundance of amino acid residues and their importance within the sequence. It is crucial to remember that the initial four positions of the peptide correspond to the N-terminal residues, while the final or last four positions correspond to the C-terminal residues. The representation of relative abundance is based on the amino acid residue with the highest significance. Upon analyzing the toxic peptides, we observe a consistent preference for the amino acid residue ‘cysteine’ across all the positions. This finding indicates that ‘cysteine’ is favored both in the N-terminal and C-terminal residues. Furthermore, at positions 1 and 3 in the C-terminus of toxic peptides, we find a higher prevalence of the amino acid residue ‘glycine’, suggesting its preference in N-terminus residues. In contrast, while examining non-toxic peptides, we discover that the amino acid residue ‘methionine’ predominantly occupies the first position. This observation implies that ‘methionine’ is the preferred residue at the beginning of non-toxic peptides. By examining these patterns of amino acid preferences, we gain valuable insights into the distinct characteristics and composition of toxic and non-toxic peptides. The implications of these findings extend our understanding of the connection between peptide structure and toxicity, paving the way for potential applications in peptide design and drug development.

**Figure 3.**
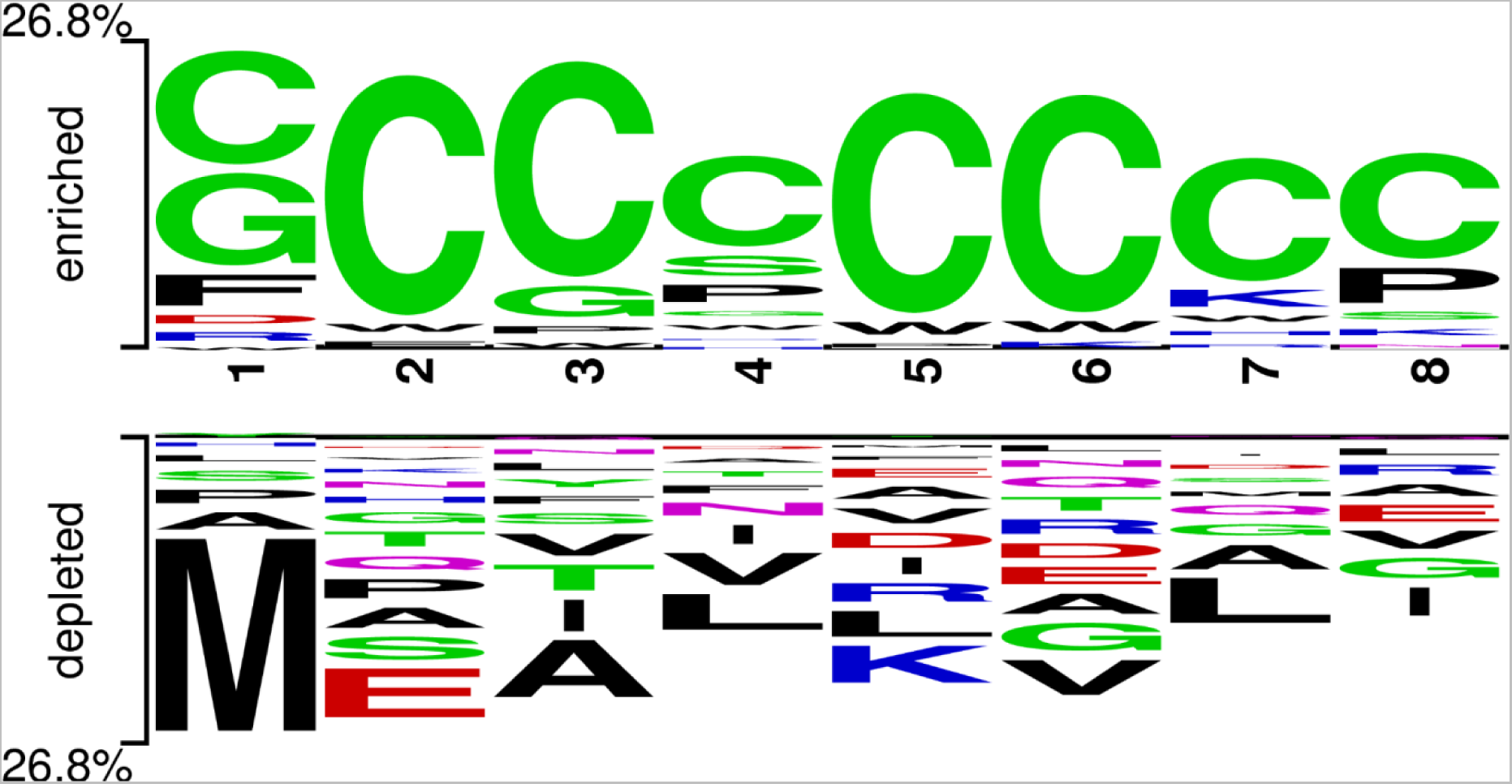
The preference of residues in toxic and non-toxic peptides at different positions is illustrated in two logo samples. First, four positions serve the N-terminus of peptides, while the last four positions serve the C-terminus of peptides.

### Similarity search using BLAST

The algorithm BLAST is often used to discover the function of a peptide or a protein on the basis of similarity search. In this study, we used BLAST to identify toxic and non-toxic peptides based on the similarity between the two sequences. We have used BLASTP-short for the e-values 10^−6^ to 10^2^. The results for BLAST are explained in detail in Table 1. It was seen that BLAST was only able to cover a part of the sequences and also had a significant error rate (I-hits). The maximum number of sequences BLAST was able to cover was 2097 out of 2208 sequences in the independent dataset with an error rate of 20.88% at e-value 10^2^. When we tried to minimize the error by decreasing the e-value, it was found that the sequences’ coverage was insufficient. At e-value 10^−6^, BLAST was able to cover only about 691 sequences out of the 2208 sequences with an error rate of 6.51%.

**Table 1:** BLAST search results for training and independent dataset.

| E-value | Training dataset |  |  |  |  | Independent dataset |  |  |  |  |
| --- | --- | --- | --- | --- | --- | --- | --- | --- | --- | --- |
|  | Total hits | Toxic |  | Non-toxic |  | Total hits | Toxic |  | Non-toxic |  |
|  |  | Chits (% of total hits) | Whits (% of total hits) | Chits (% of total hits) | Whits (% of total hits) |  | Chits (% of total hits) | Whits (% of total hits) | Chits (% of total hits) | Whits (% of total hits) |
| $10^2$ | 8366 | 3785<br>(45.24%) | 550<br>(6.57%) | 2981<br>(35.63%) | 1050<br>(12.55%) | 2097 | 913<br>(43.54%) | 166<br>(7.92%) | 747<br>(35.62%) | 271<br>(12.92%) |
| $10^1$ | 6008 | 3685<br>(61.33%) | 261<br>(4.34%) | 1745<br>(26.98%) | 317<br>(5.28%) | 1537 | 930<br>(60.51%) | 71<br>(4.62%) | 448<br>(29.15%) | 88<br>(5.73%) |
| 1 | 5241 | 3447<br>(65.77%) | 203<br>(3.87%) | 1414<br>(26.98%) | 177<br>(3.38%) | 1332 | 877<br>(65.84%) | 45<br>(3.38%) | 356<br>(26.73%) | 54<br>(4.05%) |
| $10^{-1}$ | 4752 | 3172<br>(66.75%) | 171<br>(3.60%) | 1260<br>(26.52%) | 149<br>(3.14%) | 1228 | 823<br>(67.02%) | 30<br>(2.44%) | 330<br>(26.87%) | 45<br>(3.66%) |
| $10^{-2}$ | 4269 | 2886<br>(67.34%) | 145<br>(3.40%) | 1105<br>(25.88%) | 133<br>(3.12%) | 1118 | 751<br>(67.17%) | 34<br>(3.04%) | 294<br>(26.30%) | 39<br>(3.49%) |
| $10^{-3}$ | 3827 | 2577<br>(67.34%) | 135<br>(3.53%) | 995<br>(26%) | 120<br>(3.14%) | 991 | 663<br>(66.90%) | 31<br>(3.13%) | 262<br>(26.44%) | 35<br>(3.53%) |
| $10^{-4}$ | 3394 | 2270<br>(66.88%) | 115<br>(3.39%) | 907<br>(26.72%) | 102<br>(3.01%) | 898 | 597<br>(66.48%) | 29<br>(3.23%) | 243<br>(27.06%) | 29<br>(3.23%) |
| $10^{-5}$ | 2940 | 1934<br>(65.78%) | 105<br>(3.57%) | 811<br>(27.59%) | 90<br>(3.06%) | 787 | 513<br>(65.18%) | 30<br>(3.81%) | 221<br>(28.08%) | 23<br>(2.92%) |
| $10^{-6}$ | 2569 | 1648<br>(64.15%) | 93<br>(3.62%) | 744<br>(28.96%) | 84<br>(3.27%) | 691 | 452<br>(65.41%) | 21<br>(3.04%) | 194<br>(28.08%) | 24<br>(3.47%) |

Chits = correct hits, Whits = Wrong hits

### Motif approach

The MERCI software was utilized to detect the specific patterns present exclusively in toxic and non-toxic peptides. Initially, we conducted motif searches in both the positive and negative training data. Our analysis revealed that motifs such as ‘GCCS, HPAC, CCSN, and CCSH’ were solely present in toxic peptides. We extracted 117 motifs from the positive training data and 317 motifs from the negative training data. The minimum occurrence of these motifs in both the positive and negative training datasets was 35. A complete list of motifs is provided in Supplementary Table S3. Following the motif search, we proceeded to compare these motifs against the test dataset, consisting of both toxic and non-toxic peptides. The positive motifs successfully identified only 639 peptides out of a total of 1104 in the positive test dataset. Similarly, the negative motifs were able to recognize only 353 peptides out of the total 1104 in the negative test dataset. We also conducted comparisons between the negative motifs, and the negative test dataset, as well as between the positive motifs and the positive test dataset. These comparisons resulted in 0 false positives and 1 false negative. Therefore, based on these findings, it can be concluded that the motif approach alone demonstrated poor coverage, however high probability of correct prediction.

### Deep Learning based models

In this study, we employed various deep learning models, including ANN with fixed and variable sequence length LSTM layers, ANN architecture with Bidirectional LSTM (Bi-LSTM) layers, Recurrent Neural Network (RNN) with LSTM layers and Convolutional Neural Network (CNN) with Regularized Dense layers that exercised L2 regularization. These models proved to be effective in retaining the relevant information from the training phase and adapting seamlessly to new sequences that contained diverse peptides and amino acids during the testing phase. The customized architecture of the model successfully comprehended the underlying nature of the data. The performance of all deep learning models used in the study is displayed in Table 2. ANN-LSTM with fixed sequence length achieved a maximum AUC 0.93, while ANN with Bi-LSTM layers achieved at least 0.90 AUC with balanced performance in training and testing. All the models achieved an impressive AUROC score, indicating strong predictive capability and high accuracy in distinguishing between positive and negative instances. However, it is worth highlighting that the machine learning model performed remarkably well in the classification of toxins, surpassing expectations.

**Table 2:** The performance of deep learning-based models developed using one hot encoding.

| DL | Training dataset |  |  |  |  |  | Independent dataset |  |  |  |  |  |
| --- | --- | --- | --- | --- | --- | --- | --- | --- | --- | --- | --- | --- |
|  | SENS | SPEC | PREC | ACC | MCC | AUC | SENS | SPEC | PREC | ACC | MCC | AUC |
| **ANN-LSTM- Fixed seq length** | **0.89** | **0.89** | **0.89** | **0.85** | **0.78** | **0.98** | **0.86** | **0.85** | **0.87** | **0.86** | **0.71** | **0.93** |
| **RNN with LSTM layers** | 0.87 | 0.88 | 0.88 | 0.84 | 0.77 | 0.95 | 0.86 | 0.85 | 0.86 | 0.86 | 0.72 | 0.92 |
| **CNN with Relu-Dense layers** | 0.88 | 0.88 | 0.88 | 0.84 | 0.76 | 0.95 | 0.85 | 0.85 | 0.84 | 0.84 | 0.67 | 0.92 |
| **ANN - LSTM- Variable seq length** | 0.87 | 0.87 | 0.88 | 0.84 | 0.75 | 0.95 | 0.87 | 0.94 | 0.81 | 0.83 | 0.66 | 0.91 |
| **ANN with Bi-LSTM** | 0.90 | 0.91 | 0.90 | 0.85 | 0.80 | 0.91 | 0.83 | 0.84 | 0.83 | 0.83 | 0.67 | 0.90 |

SENS = sensitivity; SPEC = specificity; ACC = accuracy; AUC = area under receiver operating characteristic; MCC = Matthews correlation coefficient

### Selected features

As discussed in the ‘Feature Selection and Ranking’ section earlier, a comprehensive evaluation of 9163 features was conducted for the datasets. In the realm of machine learning, feature selection acts as a filter, enabling algorithms to focus on the most significant aspects of the data while discarding redundant and irrelevant ones. Subsequently, after applying Tree-based feature selection methods, a total of 1373 best discriminating features were identified. Table 3 showcases the top discriminating features, which were selected not only by Tree-based feature selection but also by the majority of other feature selection methods employed in our study. AAC_C (amino acid composition of cysteine), PAAC1_C (pseudo amino acid composition of Cysteine), and QSO1_SC_C (quasi-sequence order with Schneider matrix for Cysteine) are among the best discriminatory features. Among these 1373 features Amino acid composition of cysteine (AAC_C), Quasi-sequence order with Schneider matrix for Cysteine (QSO1_SC_C), and Quasi-sequence order with Grantham matrix for Cysteine (QSO1_G_C) are the features which separately achieved 0.78 accuracy. However, when it comes to machine learning performance, we could get an increase of just 0.01 in AUC.

**Table 3:** Selected best discriminatory features in toxins and non-toxins and their corresponding accuracy and MCC using Extra Tree model.

| Features | Mean value |  | ACC | MCC |
| --- | --- | --- | --- | --- |
|  | Toxins | Non-toxins |  |  |
| AAC_C | 15.09 | 2.01 | 0.78 | 0.58 |
| QSO1_SC_C | 1.96 | 0.29 | 0.78 | 0.58 |
| QSO1_G_C | 1.02E-04 | 1.37E-05 | 0.77 | 0.56 |
| SER_C | 0.32 | 0.06 | 0.77 | 0.54 |
| ATC_S | 0.95 | 0.25 | 0.76 | 0.53 |
| PCP_SC | 0.16 | 0.05 | 0.75 | 0.52 |
| RRI_C | 0.92 | 0.23 | 0.74 | 0.49 |
| DPC1_CC | 3.47 | 0.06 | 0.74 | 0.55 |
| SEP_SC | 0.55 | 0.22 | 0.73 | 0.46 |
| QSO1_SC_C | 13640.03 | 9990.19 | 0.72 | 0.45 |

(AAC\_C = Amino acid composition of Cysteine; QSO1\_SC\_C = Quasi-sequence order with Schneider matrix for Cysteine; QSO1\_G\_C = Quasi-sequence order with Grantham matrix for Cysteine; SER\_C = Shannon entropy of Cysteine;0 ATC\_S = Atomic Composition of Sulphur; PCP\_SC = Composition of residues having sulfur content; RRI\_C = Residue repeat index of Cysteine; DPC1\_CC = Composition of Cysteine-Cysteine; SEP\_SC = Shannon entropy of residues having sulfur content, SOC1\_G1 = Sequence order coupling number with Grantham matrix for lag 1 )

### Composition-based features

To establish machine learning models, features such as AAC (amino acid composition) and DPC (dipeptide composition) were computed for both toxins and non-toxins. Among the models evaluated, the ET (Extra Trees) models exhibited outstanding performance on our dataset. This achieved a maximum AUC (Area Under the Curve) of 0.94 on the training dataset and 0.95 on the validation dataset. The performance of the dataset for composition-based features is presented in Table 5, while the performance for other features can be found in Supplementary Table S4.

### Machine learning-based models

We create prediction models using a variety of classifiers, including RF, LR, DT, ET, MLP, XGB, and SVM. The Pfeature compositional-based module was used to compute the features of the toxin and non-toxin peptides first. Using Pfeature, a total of 9163 features were calculated for each protein. We used various feature selection techniques to choose the most pertinent features, including SVC with L1 regularization, tree-based feature selection, K-best features based on statistical measures, RFE, mRMR, and VTh. The performance of the extra tree model using various feature selection techniques is displayed in the table. Among all the machine learning algorithms, tree-based classifier performed the best. It is evident that feature selection did not significantly improve performance after carefully analyzing how the ET model performed using different feature selection techniques and comparing them with results obtained without feature selection. As the majority of the features after feature selection belonged to the AAC and DPC category, we also trained our models using only the AAC and DPC compositional features, the results for which are shown in Table 4. While using the features selected by the tree-based feature selection method, the time taken for generating the results was about 315.22s, while the time taken dropped to 13.58s when we used AAC+DPC features for the model. The model achieved similar performance without a significant loss in metrics while becoming time-efficient at the same time. Also in this case, ET managed to outperform other models with an AUROC of 0.94 and 0.95 in training and testing, respectively.

**Table 4.** The performance of different feature selection methods on the ET model.

| Feature selection method | No of feature | Training dataset |  |  |  |  |  | Independent dataset |  |  |  |  |  |
| --- | --- | --- | --- | --- | --- | --- | --- | --- | --- | --- | --- | --- | --- |
|  |  | SENS | SPEC | PREC | ACC | MCC | AUC | SENS | SPEC | PREC | ACC | MCC | AUC |
| **Tree-Based** | 1373 | 0.84 | 0.90 | 0.90 | 0.87 | 0.74 | 0.95 | 0.87 | 0.91 | 0.91 | 0.89 | 0.78 | 0.96 |
| **mRMR** | 4000 | 0.83 | 0.92 | 0.91 | 0.88 | 0.75 | 0.94 | 0.87 | 0.92 | 0.92 | 0.90 | 0.77 | 0.95 |
| **VTh** | 504 | 0.84 | 0.91 | 0.91 | 0.88 | 0.75 | 0.94 | 0.86 | 0.92 | 0.92 | 0.89 | 0.76 | 0.95 |
| **SVC-L1** | 519 | 0.84 | 0.91 | 0.91 | 0.88 | 0.76 | 0.95 | 0.85 | 0.91 | 0.90 | 0.88 | 0.77 | 0.95 |
| **RFE** | 4594 | 0.85 | 0.92 | 0.91 | 0.88 | 0.77 | 0.95 | 0.84 | 0.93 | 0.92 | 0.88 | 0.76 | 0.94 |
| **K- best** | 50 | 0.80 | 0.89 | 0.88 | 0.85 | 0.70 | 0.92 | 0.78 | 0.90 | 0.89 | 0.84 | 0.68 | 0.92 |
| **No Feature Selection** | 9163 | 0.84 | 0.92 | 0.91 | 0.88 | 0.76 | 0.94 | 0.86 | 0.92 | 0.91 | 0.89 | 0.78 | 0.95 |
| **AAC + DPC** | 420 | 0.85 | 0.91 | 0.91 | 0.88 | 0.75 | 0.94 | 0.87 | 0.91 | 0.91 | 0.89 | 0.78 | 0.95 |

**Table 5.** The performance of machine learning-based models developed using AAC and DPC.

| ML | Training dataset |  |  |  |  |  | Independent dataset |  |  |  |  |  |
| --- | --- | --- | --- | --- | --- | --- | --- | --- | --- | --- | --- | --- |
|  | SENS | SPEC | PREC | ACC | MCC | AUC | SENS | SPEC | PREC | ACC | MCC | AUC |
| **ET** | **0.84** | **0.91** | **0.91** | **0.88** | **0.75** | **0.94** | **0.87** | **0.91** | **0.91** | **0.89** | **0.78** | **0.95** |
| **RF** | 0.84 | 0.91 | 0.90 | 0.87 | 0.75 | 0.94 | 0.86 | 0.91 | 0.91 | 0.88 | 0.77 | 0.95 |
| **XGB** | 0.84 | 0.89 | 0.88 | 0.86 | 0.73 | 0.93 | 0.85 | 0.89 | 0.89 | 0.87 | 0.75 | 0.94 |
| **MLPC** | 0.86 | 0.86 | 0.86 | 0.86 | 0.72 | 0.93 | 0.87 | 0.86 | 0.86 | 0.86 | 0.72 | 0.93 |
| **GB** | 0.79 | 0.88 | 0.87 | 0.84 | 0.68 | 0.92 | 0.81 | 0.89 | 0.88 | 0.85 | 0.70 | 0.92 |
| **LR** | 0.82 | 0.85 | 0.84 | 0.83 | 0.66 | 0.91 | 0.81 | 0.85 | 0.84 | 0.83 | 0.66 | 0.90 |
| **SVC** | 0.82 | 0.84 | 0.83 | 0.83 | 0.66 | 0.90 | 0.83 | 0.83 | 0.83 | 0.83 | 0.66 | 0.90 |
| **DT** | 0.81 | 0.81 | 0.81 | 0.81 | 0.62 | 0.81 | 0.82 | 0.80 | 0.81 | 0.81 | 0.62 | 0.81 |

SENS = sensitivity; SPEC = specificity; ACC = accuracy; AUC = area under receiver operating characteristic; MCC = Matthews correlation coefficient

### Machine learning combined with motif approach

To enhance the predictive capability of our model, we employed a hybrid approach in this study. This approach utilized a weighted scoring method that combined two distinct techniques: (i) a motif-based approach utilizing MERCI and (ii) a machine learning (ML)-based technique. Initially, the peptide sequence provided was classified using MERCI. Positive predictions corresponding to toxic peptides were assigned a weight of +0.5, negative predictions for non-toxic peptides were assigned a weight of −0.5, and no hits were assigned a weight of 0. Composition-based models, specifically AAC (amino acid composition) and DPC (dipeptide composition), were then integrated with this approach using various ML techniques. Among these techniques, the ET (Extra Trees) based model demonstrated exceptional performance on our independent dataset, achieving an AUC of 0.98. The performance of the combined approach of ML and MERCI for the independent dataset is shown in Table 6.

**Table 6:** The performance of the motif-based approach on the independent dataset, when combined with machine learning-based models, developed using AAC & DPC.

| ML | SENS | SPEC | PREC | ACC | MCC | AUC |
| --- | --- | --- | --- | --- | --- | --- |
| **ET** | **0.92** | **0.93** | **0.93** | **0.93** | **0.86** | **0.98** |
| **RF** | 0.90 | 0.94 | 0.93 | 0.92 | 0.84 | 0.98 |
| **XGB** | 0.89 | 0.94 | 0.94 | 0.91 | 0.84 | 0.98 |
| **GB** | 0.89 | 0.92 | 0.92 | 0.91 | 0.82 | 0.97 |
| **SVM** | 0.85 | 0.93 | 0.93 | 0.89 | 0.79 | 0.96 |
| **MLPC** | 0.86 | 0.94 | 0.93 | 0.90 | 0.81 | 0.96 |
| **LR** | 0.83 | 0.94 | 0.93 | 0.88 | 0.78 | 0.96 |
| **DT** | 0.82 | 0.80 | 0.80 | 0.81 | 0.63 | 0.87 |

SENS = sensitivity; SPEC = specificity; ACC = accuracy; AUC = area under receiver operating characteristic; MCC = Matthews correlation coefficient

### Deep learning combined with motif approach

In our study, we have utilized an ensemble methodology to integrate the motif-based MERCI software with an advanced deep learning model to achieve precise prediction of protein toxicity. In order to combine both the scores of DL and MERCI, we did the same procedure as in combining the ML and MERCI. This innovative strategy leveraged the strengths of both approaches, resulting in a significant improvement in the AUC. Table 7 presents the performance of the combined approach involving Deep Learning and MERCI on the independent dataset. Among the DL techniques we used, the ANN - LSTM - Fixed sequence length demonstrated the best performance on the independent dataset, achieving an AUC of 0.97. Notably, after combining the results of DL and MERCI, the performance comes very close to the ML and MERCI hybrid model.

**Table 7:** The performance of the deep learning combined MERCI-based models on the independent dataset.

| **DL** | **SENS** | **SPEC** | **PREC** | **ACC** | **MCC** | **AUC** |
| --- | --- | --- | --- | --- | --- | --- |
| **ANN - LSTM-Fixed seq length** | **0.92** | **0.91** | **0.90** | **0.92** | **0.81** | **0.97** |
| **ANN - LSTM-Variable seq length** | 0.87 | 0.95 | 0.94 | 0.91 | 0.90 | 0.96 |
| **ANN with Bi-LSTM layers** | 0.86 | 0.95 | 0.94 | 0.90 | 0.81 | 0.96 |
| **CNN with Relu-Dense Layers** | 0.85 | 0.95 | 0.95 | 0.90 | 0.80 | 0.96 |
| **RNN with LSTM layers** | 0.86 | 0.96 | 0.95 | 0.89 | 0.81 | 0.95 |

SENS = sensitivity; SPEC = specificity; ACC = accuracy; AUC = area under receiver operating characteristic; MCC = Matthews correlation coefficient

We hereby understood that neural networks perform really well in peptide sequence classification and can be used further in various vaccine or antibiotic generation processes. This research showcases the effectiveness of deep learning architecture and its future usage in bioinformatics.

### BLAST Search

To develop a robust prediction methodology, we integrated the similarity search method BLAST with separate machine learning (ML) and deep learning (DL) approaches. Initially, the BLAST search was conducted on a query sequence, and if a significant hit was found, the query sequence was classified as toxic or non-toxic based on the BLAST result. However, if the BLAST search did not yield any hits, an alternative approach, either ML or DL, was employed. We compared the performance of the ML-based approach to the DL-based approach combined with BLAST as displayed in Table 7; we observed a decrease in performance for the ML model, which decreased from AUC 0.95 to 0.938 on the independent dataset. Conversely, the DL model demonstrated an improvement in performance, with the AUC increasing from 0.90 to 0.919. We further examined the performance of a combined approach involving ML or DL, the MERCI method, and BLAST. Unfortunately, combining BLAST with ML MERCI hybrid and DL MERCI hybrid did not improve the performance. It is worth noting that only the MERCI hybrid models achieved the best performance in our evaluation.

### Implementation of ensemble approach

In this study, we developed two hybrid approaches: the first hybrid approach combined MERCI with ML techniques, while the second hybrid approach integrated MERCI with Deep Learning (DL) techniques. These two hybrid approaches were carefully designed and implemented to exploit the complementary strengths of both the involved methods. By leveraging the unique qualities of MERCI and either ML scores or DL scores, the study aimed to achieve superior performance and accuracy compared to using each approach independently. The hybrid approach has been used in several studies and has been shown to improve the accuracy of peptide classification. A comparison between the performance of both ML and DL models as well as their hybrid model, is shown using a spider graph in Figure 4.

**Figure 4:**
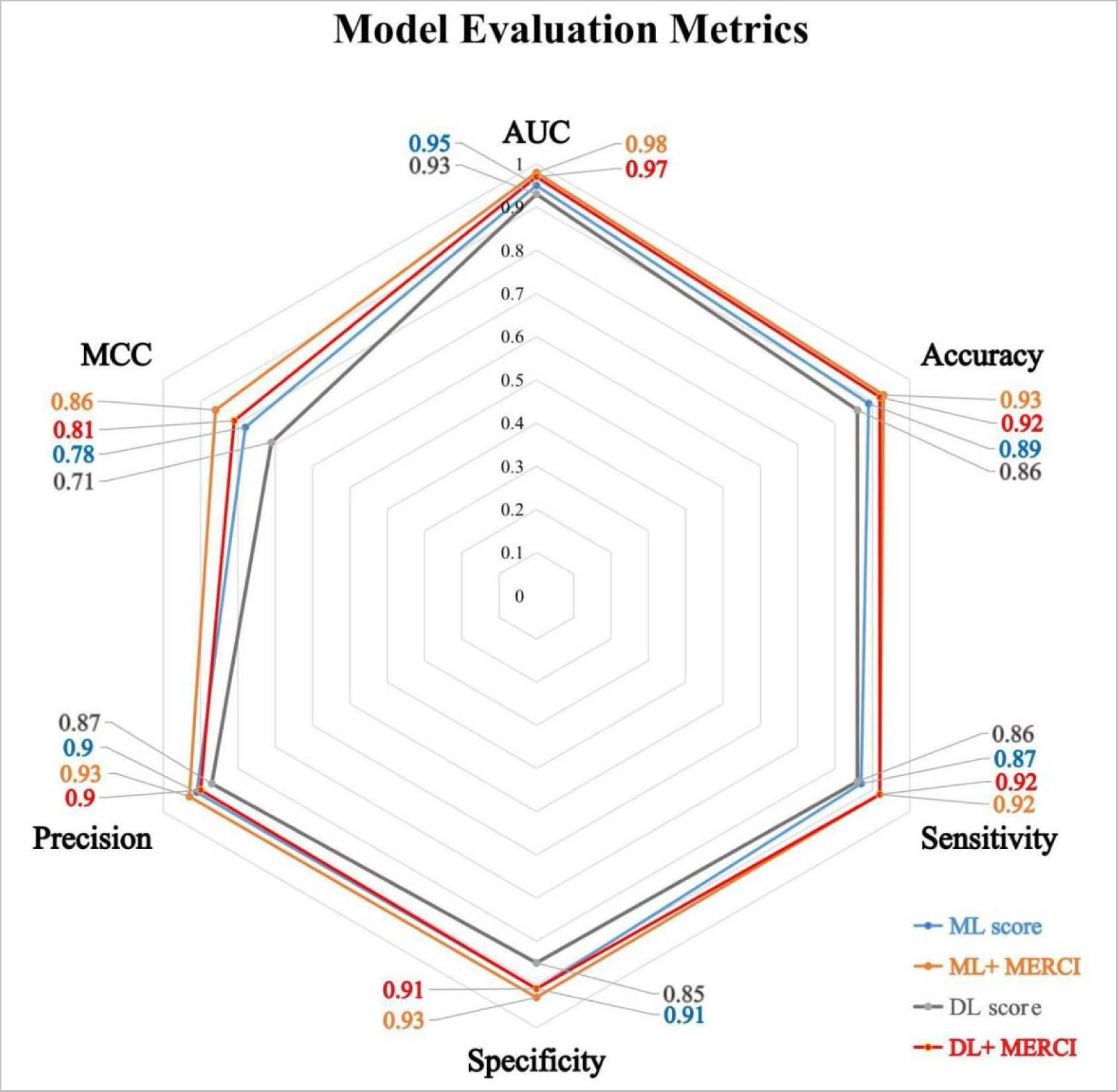
A comparative overview of the performance of machine learning, deep learning and the ensemble model (with MERCI).

### ROC Plot

We employed Receiver Operating Characteristic (ROC) curves to assess the performance of our models without relying on a specific threshold. Using the RocCurveDisplay function from scikit-learn, we generated ROC curves with the corresponding Area Under the Curve (AUC) values. As depicted in Figure 5, the hybrid of ML and MERCI method exhibited a higher AUC value than ML, DL, and the hybrid of DL and MERCI. This signifies that the ML and MERCI hybrid methods accurately classified toxic proteins.

**Figure 5:**
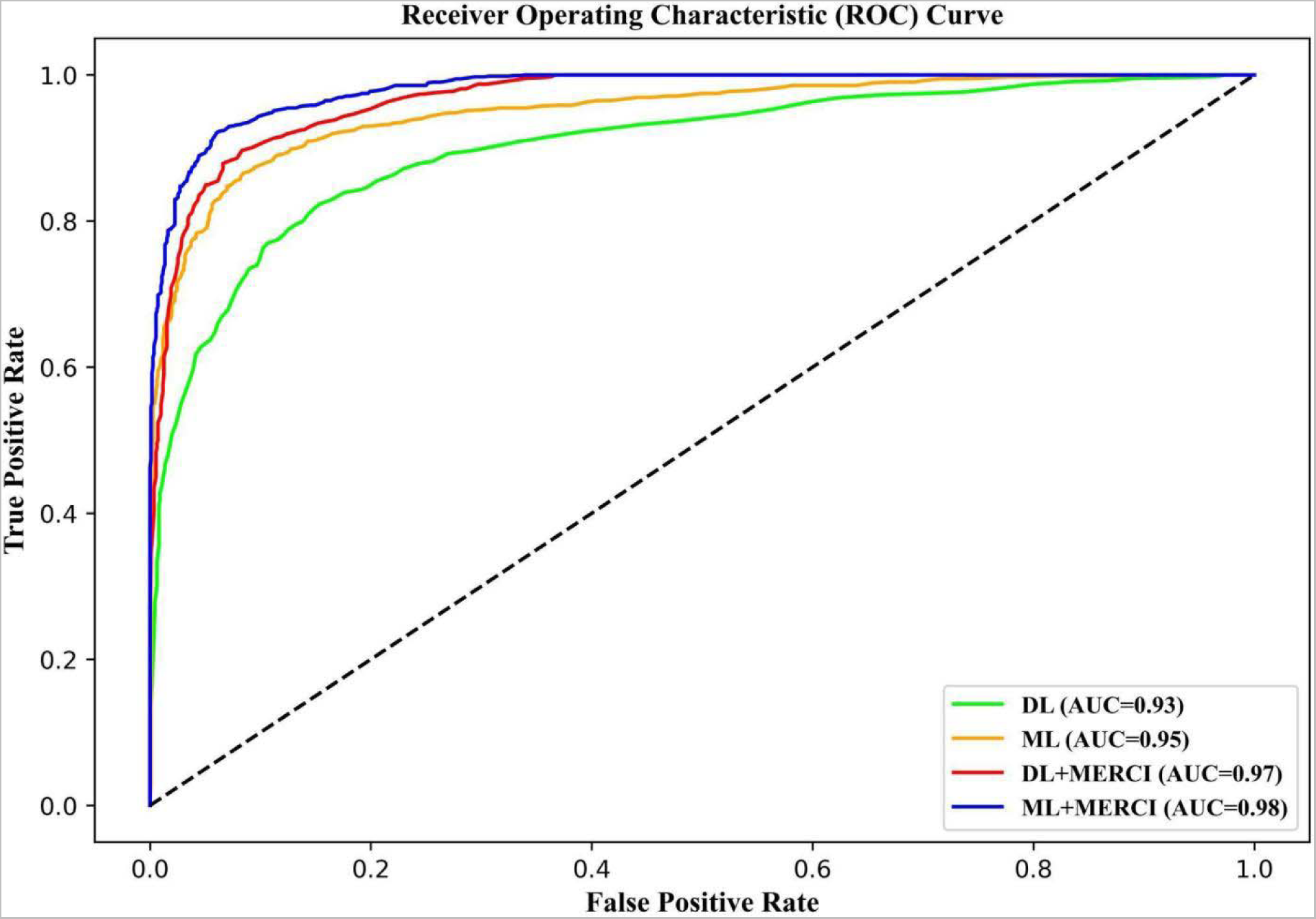
ROC curves of all the models based on machine learning and deep learning methods as well as their hybrid with MERCI.

### The development and deployment of a web server

For the purpose of predicting toxic peptides, ToxinPred3.0 (https://webs.iiitd.edu.in/raghava/toxinpred3) was created. We have implemented ML and DL models as well as their MERCI-hybrid model. The web server includes the primary modules, including (a) prediction, (b) Protein scanning, (c) motif scan, (d) BLAST search, and (e) download. The ‘prediction module’ enables users to submit single or multiple peptide sequences in FASTA format. Based on the AUC values of the peptide sequence, this module can effectively distinguish between toxic and non-toxic peptides. We have also calculated positive predictive values (PPV) for all the predictions. The positive predictive value represents the probability that a positive prediction (indicating toxicity) accurately indicates that the peptide is truly toxic. ‘Protein scanning module’ will help to identify the toxic regions in the protein. The ‘motif scan module’ locates the motifs only found in toxic peptide sequences. Additionally, it maps or scans the user-provided query peptide sequence’s motif data and classifies toxic and non-toxic motifs according to their presence or absence. The ‘BLAST search module’ is a valuable tool provided by our platform that enables users to perform similarity-based searches using BLAST (Basic Local Alignment Search Tool) against a comprehensive database of toxins and non-toxins. This search module serves as a powerful resource for users to explore and identify similar sequences within the database. To enhance user experience and accessibility, we have developed a web server with a responsive HTML template. This design ensures that users can conveniently access and navigate the platform across various operating systems and devices, providing a seamless and user-friendly experience. We have created both a standalone package and a pip package, providing users with the capability to predict toxins on a comprehensive scale. This versatile and efficient solution is ideal for extensive analyses, accessible and downloadable through our web server’s dedicated ‘Download’ module.

### Comparison with other methods

To support the creation of a new technique, it is crucial to evaluate how well the suggested method performs in comparison to already available methods. In Table 8, we compare the effectiveness of the suggested approach, ToxinPred3.0, with other currently used methods as stated in the literature. As can be seen in Table 8, ToxinPred3.0 outperforms other existing methods.

**Table 8:**
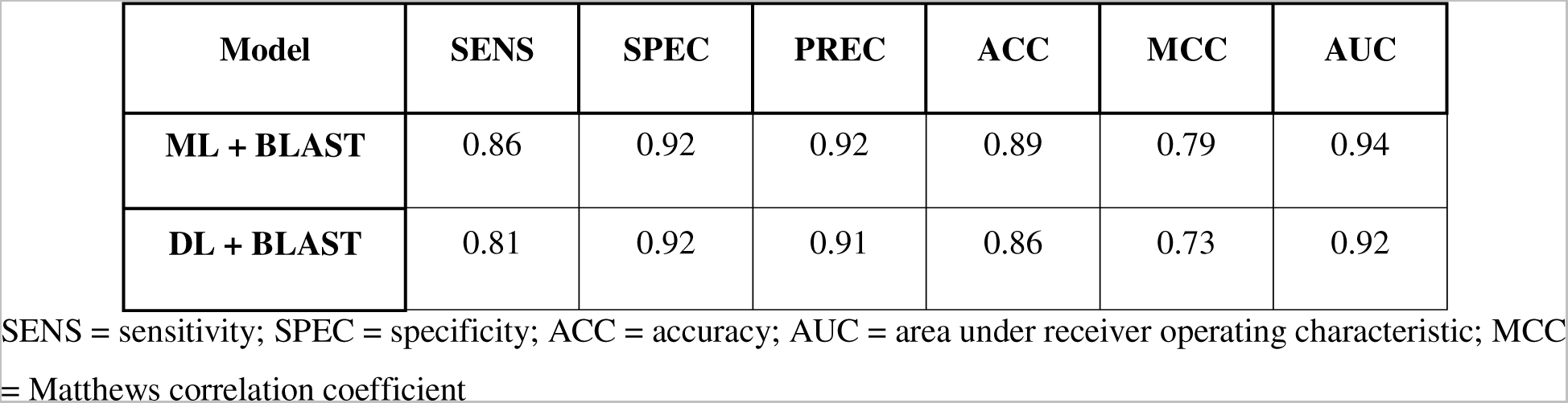
Performance of BLAST-based approach on independent dataset when combined with best performing models of ML and DL.

| Model | SENS | SPEC | PREC | ACC | MCC | AUC |
| --- | --- | --- | --- | --- | --- | --- |
| **ML + BLAST** | 0.86 | 0.92 | 0.92 | 0.89 | 0.79 | 0.94 |
| **DL + BLAST** | 0.81 | 0.92 | 0.91 | 0.86 | 0.73 | 0.92 |

SENS = sensitivity; SPEC = specificity; ACC = accuracy; AUC = area under receiver operating characteristic; MCC = Matthews correlation coefficient

We evaluated the effectiveness of current tools on the independent dataset used in our study to provide a comparison with other existing approaches. For the independent dataset utilized in ToxPred3.0, ToxinPred and ToxIBTL obtained AUC values of 0.85 and 0.80, respectively, as stated in Table 8. The long protein toxicity prediction algorithms (ToxMVA, ToxinPred2, ToxClassifer, TOXIFY, and ToxDL) cannot be used because the dataset used in ToxinPred3.0 contains only peptide/ short proteins sequences. Moreover, we were unable to predict the toxicity of proteins in the independent dataset of ToxinPred3 using ATSE and CNN_BiGRU due to the limitations of their services (ATSE - improper web service functioning, and CNN_BiGRU-standalone is not functional).

The significance of the recently suggested strategy in the area of toxicity prediction is shown by this comparison in Table 8.

**Table 9.**
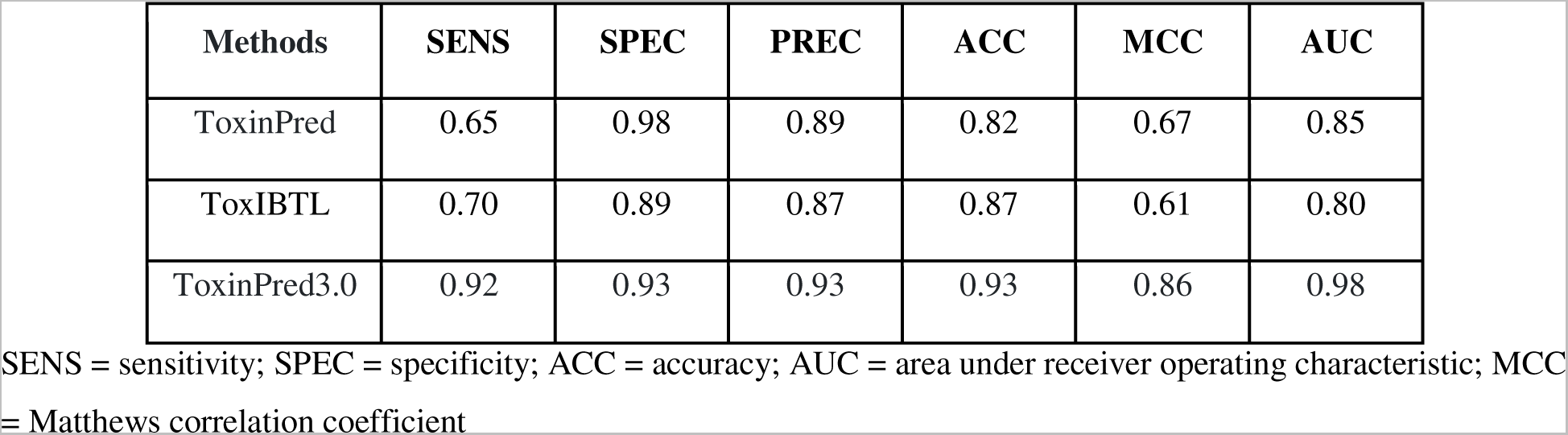
Comparison of Performance of existing methods on independent dataset of ToxinPred3.0.

## Discussion

A crucial step in the development of peptide-based drugs is protein toxicity prediction. It is possible to shorten the duration and expense of experimental testing by predicting protein toxicity reliably [11]. The current work aimed to create an in-silico model that could anticipate the toxicity of therapeutic peptides with very high accuracy.

The dataset used in this investigation included 5518 known toxic peptides from several databases as well as 5518 non-toxic peptides that were only gathered from SwissProt. These peptides were thoroughly analyzed using a variety of strategies, including sequence alignment, motif recognition, machine learning, and deep learning techniques.

The Pfeature tool was employed to calculate a range of features for all the peptide sequences. Initially, a vector consisting of 9163 features was computed for each sequence in the dataset. Our high-performing model was assessed using three sets of features: the entire feature set, AAC and DPC-based composition features, and a subset obtained from the extensive feature pool through feature selection. To accomplish this, we employed various feature selection techniques, which aimed to identify a smaller subset of pertinent features from the original set, while eliminating irrelevant, redundant, or noisy features. Despite employing various feature selection methods to identify and rank relevant characteristics, the performance of the model based on these specified features did not exhibit a significant improvement. Notably, composition-based features were consistently selected as important features, as they effectively distinguished between toxic and non-toxic peptides.

Our preliminary analysis of toxic peptides showed that Cys was highly abundant and preferred at most positions in toxic peptides, as determined by calculating the percent amino acid composition. Additionally, the composition of Pro, Asp, and His was found to be considerably higher in toxic peptides compared to non-toxic peptides. Dipeptide analysis, which considers the pairwise combination of amino acids, has shown superior performance over composition-based models in previous studies. In this study, we aimed to improve peptide toxicity prediction by incorporating both AAC and DPC information. To achieve this, binary profiles were generated, and machine learning (ML) models were developed using these profiles as input features. The models based on AAC combined with DPC-based features outperformed the models solely based on simple amino acid composition, demonstrating the importance of considering both composition and amino acid order in predicting peptide toxicity. Other than ML, we also implemented deep learning (DL) models based on simple binary profiles. However, ML performed better compared to DL.

Additionally, we have incorporated the use of BLAST, a commonly employed tool for annotating peptide sequences. By comparing the query peptide sequence to a vast database of known peptide functions, BLAST can assign the same function to the query peptide if a significant similarity is identified. We performed BLAST-shortp to determine whether a query peptide is toxic or non-toxic. The algorithm was implemented for e-values ranging from 10^−6^ to 10^2^. It was observed that at e-value 10^2^, BLAST was able to cover a large number of sequences but with a high error rate of 20.84% on the independent dataset, whereas, at the e-value of 10^−6^, it was able to cover a low number of sequences with an error rate of about 6.5%. BLAST alone was not able to distinguish between toxic and non-toxic peptides with high accuracy, and after combining the scores from BLAST with Deep Learning and Machine Learning based methods, it was discovered that the overall accuracy was constant or decreasing because of the incorrect identifications by BLAST (~2.14%). Hence, we decided to exclude BLAST from the final hybrid models.

In our study, we have harnessed the power of deep learning, a prominent technique widely employed in natural language-related tasks, to tackle the challenging problem of toxin prediction. Leveraging the versatility of deep learning architectures, we explored several existing models designed for sequence classification. Ultimately, we found that a custom-built artificial neural network incorporating LSTM layers, implemented using the Keras API, delivered the most promising performance. To mitigate overfitting concerns, we applied L2 regularization to ensure the generalizability and robustness of the model. In this case, we achieved a maximum AUC score of 0.93. Notably, the performance of the machine learning model was superior to deep learning. However, the AUC increased to 0.97 when we merged the deep learning and MERCI scores, which is practically as good as our final ML and MERCI hybrid model.

Furthermore, we created a comprehensive platform that facilitates the classification of proteins and peptides into toxic and non-toxic categories. To ensure widespread accessibility, we have developed a web server and a standalone package for ToxinPred3.0. The web server offers a user-friendly interface and incorporates the most effective model for accurately predicting toxins and non-toxins. Our standalone package allows users to utilize the prediction method offline, granting flexibility in their analyses. While our model exhibits promising performance, we acknowledge the need for further enhancements. Continuous research and refinement of the prediction method will contribute to improved accuracy and reliability. We encourage researchers to employ our prediction method in their work, leveraging it to design enhanced protein and peptide-based therapeutics targeting diverse diseases. By providing this comprehensive platform, we aim to advance the protein and peptide therapeutics field, empowering scientists to make informed decisions regarding the toxicological aspects of their molecules.

## Authors’ contributions

ASR and AA collected and processed the datasets. ASR and PT implemented the algorithms and developed the prediction models. GPSR analyzed the results. SC created the back end of the web server, and SC and ASR created the front-end user interface. ASR, AA, PT, and GPSR penned the manuscript. GPSR conceived and coordinated the project. All authors have read and approved the final manuscript.

## Acknowledgments

The authors express their gratitude to the University Grants Commission (UGC), Council of Scientific and Industrial Research (CSIR), and Department of Science & Technology (DST) for their generous fellowships and financial support. They also thank the Department of Computational Biology, IIITD, New Delhi, for the excellent infrastructure and facilities. The authors would like to acknowledge the Department of Biotechnology (DBT) for the infrastructure grant awarded to the institute. Furthermore, they would like to acknowledge BioRender.com for the creation of the figures utilized in this work.

## Data Availability Statement

The datasets generated for this study can be accessed on the ‘ToxinPred3.0’ web server at https://webs.iiitd.edu.in/raghava/toxinpred3/download.php. The source code for this study is publicly available on GitHub and can be found at https://github.com/raghavagps/toxinpred3.

